# Vertebrates but not ants protect rainforest trees from herbivorous insects along an elevational gradient in Papua New Guinea

**DOI:** 10.1101/2022.07.03.497915

**Authors:** Katerina Sam, Leonardo Re Jorge, Bonny Koane, Pita Amick, Elise Sivault

## Abstract

**Aim:** The theory on trophic interactions between plants, insect herbivores, and their predators predicts that predators increase plant biomass by feeding on herbivores. However, it remains unclear whether different types of predators regulate herbivores to the same degree, and how the trophic interactions affect lower trophic levels along elevational gradients where predator communities differ significantly. Therefore, we aimed to investigate the impact of vertebrate predators and ants (individually and in combination) on arthropod communities and leaf herbivory along a complete tropical forest gradient.

**Location:** Papua New Guinea

**Taxon:** Multi-taxon

**Methods:** We excluded predators from 560 saplings in two six-month long predator exclusion experiments spanning wet and dry seasons. Saplings were spread across 8 study sites which were evenly spaced at 500 m elevational increments from 200 to 3700m a.s.l..

**Results:** On average the density of arthropods increased significantly by 37% and 33% respectively when vertebrate predators, and both ants and vertebrates predators, were removed. Both season and elevation mediated this effect significantly. At lower trophic levels, both the exclusion of both vertebrates alone, and exclusion of vertebrates plus ants, led to a significant increase in leaf damage by 50% and 36% respectively. In contrast, the exclusion of ants alone had no significant effect on arthropod density or leaf damage, which increased by 12% and 9% respectively.

**Main conclusions:** Our results indicate that the relative contribution of birds and bats changes at different elevational sites, while the overall effect of vertebrate predators remains consistent along the whole elevational gradient. This contrasts with ant driven trophic cascades which brought about increased herbivory only at the most productive sites of the elevational gradient, where ant abundance is highest. We conclude that disappearance of insectivorous vertebrate predators can lead to substantial negative consequences for plants.

## Introduction

Insect herbivores are the primary consumers of plant leaf tissue in tropical forests (Coley and Aide 1991, Coley and Barone 1996, but Suzuki, et al. 2013), and their impact is notably increased in the absence of top-down control (Mooney, et al. 2010, Sam, et al. 2022). This top-down control is therefore a high value ecological service across a range of tropical ecosystems (Schmitz 2006, Sekercioglu 2006). It is usually assumed that disparate insectivorous predator groups have similar effects and are thus treated as a single functional unit, leaving their individual or joint effects largely unknown (Mooney 2007, Perfecto and Vandermeer 1996, Richards and Coley 2007, Sih, et al. 1998). Resolving the often-complex trophic dynamics of multiple types of predators in the same community and their cascading effect on plant performance remains an important challenge in ecology (Singer, et al. 2017). This challenge is primarily rooted in the variable strength of trophic cascades between predators, prey and plants both within (Moon and Stiling 2004, Mooney and Linhart 2006) and between communities (Shurin, et al. 2002).

Bats and birds, and to a lesser extent insectivorous vertebrate, consume large quantities of insects and other arthropods (Kalka and Kalko 2006, Mooney, et al. 2010, Nyffeler, et al. 2018). They are likely to act as intraguild predators, feeding both on predatory as well as herbivorous arthropods (Ingala, et al. 2021, Karp and Daily 2014, Milne, et al. 2016, Sam, et al. 2017). The impact of ubiquitous insectivorous birds on lower trophic levels was shown in some but not all studies (Mooney, et al. 2010, Van Bael, et al. 2008). The impact of bats on arthropods and herbivory might be comparable to the effect of birds according to some studies (Kalka and Kalko 2006, Williams-Guillén, et al. 2008). However, bats can be erroneously overlooked in exclusion studies (Mooney, et al. 2010, Sam, et al. 2022), as the typical approach of covering foliage with a net effectively quantifies the combined effect of excluding both birds and bats (Greenberg, et al. 2000, Holmes, et al. 1979, Van Bael, et al. 2003, but see Kalka, et al. 2008).

Ants are recognised as other important predators in many habitats (Hölldobler and Wilson 1990, Rosumek, et al. 2009). Despite their study as natural enemies and biological control agents (Mestre, et al. 2012, Philpott and Armbrecht 2006, Rosumek, et al. 2009, Way and Khoo 1992), their importance as predators remains equivocal. Previous efforts failed to detect their effect on lower trophic levels or showed temporal inconsistency (Tanhuanpää, et al. 2001, Sam, et al. 2022). Plant damage in the absence of ants was reported to increase between 0 and 250% on average, with the duration of the ant exclosure (Sam, et al. 2022), and productivity of the study site (Rosumek, et al. 2009, Sam, et al. 2022) having signficant impacts. Ant exclosures are difficult to conduct (Hood, et al. 2022, Rosumek, et al. 2009) thus bait removal is often considered as a proxy of predation (e.g., Liu, et al. 2020, Roslin, et al. 2017, Burger, et al. 2021). Unfortunately, this method may not accurately measure the realized top-down control and trophic cascade to plants from ants.

Changes in the diversity and abundance of animal taxa along elevational gradients is one of the most striking biogeographic patterns on Earth (Colwell, et al. 2016, McCain 2005, Rahbek 1995, Sam, et al. 2019). These distributional changes lead to changes in the relative importance of predator groups at different sites on the same gradient. As such, elevational gradients provide an excellent experimental environment for studies focussing on the individual and combined effects of insectivorous predators on prey and plants (Schemske, et al. 2009). As climate change is expected to lead to further changes in predator distribution and thus ecosystem function (Chen, et al. 2011), it is increasingly important to understand individual and combined effects of various groups of predators along large environmental gradients.

Changes in diversity along elevational gradients are also accompanied by changes in body size where lowland animals are generally larger than animals at higher elevational sites. As predator size also determines preferred prey sizes, we can also expect the effect of predators to be detectable in the mean body size of insect communities. For example, birds were shown to reduce the mean size of caterpillars by 12% through preferentially feeding on large individuals (Singer, et al. 2017). Similarly, birds and bats reduced mean body size of Araneae, Formicidae and arthropod larvae (Karp and Daily 2014). In contrast to vertebrate predators, ants are expected to hunt smaller prey, thus their absence from an ecosystem might lead to an increase in mean arthropod body size.

Here we investigated the impact of vertebrate predators and ants (individually and in combinations) on arthropod communities and leaf herbivory along a complete tropical forest gradient in Papua New Guinea. We aimed to answer the following questions: (1) Do predators significantly affect the density of arthropods in various tropical habitats along the elevational gradient? We hypothesise that the removal of vertebrate predators, but not ants, will result in an increased density of arthropods on plants. Moreover, we expect that the effect of predators will decrease with increasing elevation (1A) and will be higher in the wet season than in the dry season (1B) due to the differences in productivity and the assumption that tropic interactions are stronger in more productive environments. (2) Are the effect of ants and vertebrate predators detectable on the mean body size of arthropod communities? We propose that arthropods will be relatively larger in the absence of vertebrate predators and relatively smaller in the absence of ants. (3) Do any of the predator groups display disproportionate impacts on predatory arthropod abundance compared with herbivore abundance? We expect that both focal predator groups (i.e., vertebrate predators and ants) act as intraguild predators whereby they feed on both herbivorous and predatory arthropods similarly, thus diminishing the effect of trophic cascades but not so much as it wouldn’t be detectable. (4) Are the effects of ants and vertebrate predators independent of one another or do they interact? We hypothesise that the effect will be mostly independent and additive, as ants and vertebrate predators feed on different prey items and birds (and presumably also bats) consume few ants (Sam, et al. 2017).(5) Does the potential effect of predators cascade down to the trophic level of plants and is it detectable on plant herbivory? Considering our first hypothesis, we suggest that the removal of vertebrate predators, but not ants, will have a measurable effect on plants thus resulting in increased herbivory.

## Materials and methods

### Study sites and experimental trees

Our study was performed along the slopes of Mt. Wilhelm (4,509 m a.s.l.) in the Central Range of Papua New Guinea (Figure S1). The rainforest gradient spans from the lowland floodplains of the Ramu river (200 m a.s.l., 5° 44’ S 145° 20’ E) to the timberline (3,700 m a.s.l., 5° 47’ S 145° 03’ E). The gradient is 30 km long and consists of eight sites evenly spaced at 500 m elevational increments. Average annual precipitation is 3,288 mm (local meteorological station) in the lowlands, rising to 4,400 mm at 3,700 m a.s.l., with a distinct condensation zone around 2500 – 2700 m a.s.l. Mean annual temperature (measured by our data loggers – Sam, et al. 2019) decreases from 27.4°C at the lowland site to 8.4°C at the tree line at a constant rate of 0.54°C per 100 elevational metres. The habitats and zonation of forest types are described elsewhere (McAlpine, et al. 1983, Paijmans 1976, Sam, et al. 2019, Tvardikova 2013).

We selected and tagged saplings belonging to 3 – 7 tree species at each elevational study site prior to the experiment (Table S1). Unfortunately, no single tree species or genus is distributed along the complete elevational gradient. Therefore, we aimed to work with several *Ficus* species which represent the most dominant and ecologically important genus from 200 up to 2700 m a.s.l. We had to work with other locally dominant species at the two highest elevations (Table S1). In total, we selected 80 (or 40 at 3,200 and 3,700m asl) individual saplings per elevational site, i.e., 560 saplings along the entirety of the elevational gradient (Table S2). For statistical independence, we allowed at least 50 m between any pair of tree individuals. We visually assessed saplings of the focal species and selected individuals which had approximately 500 leaves growing within a well-developed crown 2.5 – 4 m above the ground, did not have any ant nests, and did not have abnormally high herbivory or fungal damage. The selected tree species did not produce any exudates or sugar droplets for attracting ants. Average leaf-sizes of the selected species ranged from 16.31 to 154.10 cm^2^ (mean ± S.E. = 52.16 ± 6.12). Mean sapling DBH at the beginning of the experiment was 1.16 cm for exclosures and 1.17 cm for controls.

### Experimental design

We prevented vertebrate predators (i.e., birds and bats together, VER) from accessing 20 out of the 80 preselected saplings at each elevational site between 200 and 2,700 m a.s.l. We also prevented ants accessing another 20 saplings (ANT) and both vertebrate predators and ants accessing another 20 saplings (ALL) at each elevation. We left 20 saplings without predator protection as controls (CON). At the elevational sites 3,200 and 3,700 m, we protected 20 saplings against vertebrates (VER) and kept 20 as control saplings (CON). There were no ant exclosures at these two sites as they are above the natural elevation range of ants (Moses 2015, Colwell et al. 2016). Thus, at each elevation, 3-7 tree species were selected for conducting our experiments, with a total of 18 tree species used across the 6 elevation bands between 200 m and 3700 m. Most tree species occurred in multiple elevation bands (see Table S1 for details).

Vertebrate (VER) exclosures were constructed from PVC tubes (1 cm in diameter) joined together by PVC corners and covered with agricultural nylon netting (mesh opening 3 × 3 cm, transparent green). Each exclosure had dimensions 2 × 2 × 2.5 m and comprised a volume of 10 m^3^, which enclosed an average of 1.63 (± S.D. 1.02) m^2^ of leaf area. The mesh size was comparable to other exclosure studies [e.g., 29×29 mm in Greenberg, et al. (2000); 25×25 mm in Mols and Visser (2002); 20×20 mm in Van Bael, et al. (2003)]. The exclosure materials did not attract arthropods, did not damage leaves or branches, and did not significantly reduce light. We attached a system of ropes to neighbouring vegetation (i.e., strong branches) in order to secure the exclosures in place (thus preventing the exclosure from moving and disturbing the foliage of the target saplings). Observations confirmed that netting did not exclude small lizards as they were seen crawling under the cage. However, one ca. 50 cm long lizard was found tangled in the netting and was subsequently released, suggesting the exclosures partially prevented access of non-flying insectivores which were not quantified in the study.

To exclude ants (ANT), we used the adhesive Tanglefoot® pest barrier (Philpott, et al. 2008, Philpott, et al. 2004) which we applied in a 30cm wide strip around the trunk of the sapling at breast height. We removed all lianas and branches touching the sapling and we carefully checked for potential ant nests in plant hollows. The ants foraging on foliage were removed by both beating and manual removal. We placed tuna baits (one teaspoon under a gauze tied to the trunk of the tree) 20 cm above the Tanglefoot layer and checked for the presence of ants two hours later. All individuals were removed. Tanglefoot was reapplied every 3 – 4 weeks and new tuna bait was placed at the same place to confirm that ants were not present. Some ants crossed the Tanglefoot barriers so our analysis excluded all ant exclosure saplings where ant activity exceeded 3 ants on a single checking date (3 saplings in total).

To exclude all (ALL) predators (ants and vertebrate predators) we used a combination of the above methods, i.e., by applying Tanglefoot to the trunk, installing a cage around the sapling, and preventing the foliage from touching the cage. Tanglefoot was also applied to all of the support ropes used to lower the cage and hold it in place so as to prevent ants from accessing the sapling via neighbouring vegetation.

### Measured variables

We sampled the effect of predator exclusion after 6 and 12 months, corresponding to the dry and wet season respectively (Table S2). We first carefully opened the protective nettings where needed. Next the trunk of the sapling was slowly lowered above a 2×2m mosquito net, wrapped into the net, and sprayed by fast knock-down insecticide (Mortein®). Shortly after we shook the foliage firmly, opened the net, collected all arthropods (>1mm) and preserved them in vials filled with DNA grade ethanol. Arthropods were later identified into orders or families, feeding specialization, life stage and measured to the nearest 0.1 mm (Table S3).

To assess herbivory leaf damage, we randomly selected two branches (ca. 50 leaves in total) and collected all their leaves. We estimated how many other leaves in total were present on the sapling and used this number to calculate total leaf area. We took photographs of the flattened leaves of each sapling against a 50 × 50 cm^2^ white background. Using Adobe Photoshop CS6 (Adobe Systems Inc., USA), we outlined the missing edges of the leaves based on their expected shape. We then used ImageJ version 1.47 (National Institute of Health, USA) to calculate the remaining leaf area (a, in cm^2^), the extrapolated leaf area without any herbivore damage (b), and the area lost to herbivory (c = b - a). We then estimated the percentage of leaf–area loss per m^2^ of foliage and total standing leaf area per each whole focal sapling using the count of the number of leaves (i.e., number of leaves * mean leaf area a) which we later used to calculate densities of arthropods per m^2^.

We completed the second survey on the same saplings approximately six months after the first survey (Table S2). After the first survey, all exclosures were renewed and only some (n= 16) saplings (including their treatments) were replaced completely due to their death or damage (e.g., vandalism of villagers, fallen branches, landslides). The second survey was conducted exactly the same way as the first, with the only difference being that *all* foliage was now collected from each sapling. We placed all leaves collected from individual saplings into a bag, randomly selected ca. 50 leaves for photographing and further analyses, and weighed all leftover foliage as well as the leaves used for photography. This allowed us to calculate total leaf area (from known weight and leaf area of leaves used to measure herbivorous damage) of the saplings more accurately than during the first survey. We found that leaf herbivory within a branch (i.e., herbivory on leaves from first survey) was correlated similarly (P < 0.001, R = 0.854) to herbivory on leaves randomly collected within the whole plant (i.e., herbivory on leaves from first survey; P < 0.001, R = 0.628). Further, estimated leaf area (from the first survey) was correlated (P < 0.001, R = 0.358) to the real leaf area based on leaf-weight (from the second survey), despite some saplings growing significantly more than others during the project.

### Predator survey

Bird communities here have been repeatedly surveyed by point counts as part of long-term monitoring efforts, with this methodology and data published in several studies. In short, point counts at each elevational site were carried out at 16 points regularly spaced along a 2,350 m transect (successive points were 150 ± 5 m apart to avoid overlap). All birds seen or heard within a fixed radial distance of 0 - 50 m were recorded. Each count lasted 15 min. Bat communities were surveyed in the understory during two expeditions conducted in wet (February – March 2015) and dry seasons (June – July 2015). This method and data were published in Sivault, et al. (2022). In short, we used an ultrasonic bat call detector coupled with a recorder to detect echolocating bat species. We recorded bats at five points (i.e., 15 minutes per point) separated by 200 meters at each elevation, in line with the bird transect methods described above. Surveys were conducted for four days per site after sunset (6 pm). **T**he ant communities at each of the study sites were sampled by hand collection. The trunk of each sapling was inspected for ants at breast-height for 10 minutes, and also by the tuna bait method in May-June 2014. More details on the respective survey are provided in Supplementary material.

## Data analysis

### Arthropod densities

We used a linear mixed model and model selection to determine the effect of our experimental treatments on arthropod densities. We used arthropod density (per m^2^ of foliage) as our response variable and treatment, season and elevation and possible two-way interactions between them as our predictor variables. Elevation was modelled as a second-order polynomial to allow for non-linear elevational trends. We used tree species as a random effect in our model to account for differences in baseline arthropod densities between species, an expectation arising from variation in defensive traits. Individual saplings were used as a second random factor as each of the 560 focal saplings was surveyed twice i.e., in both wet and dry seasons.

### Herbivory

*The* effect of our experimental treatments on herbivory damage (as the proportion of the leaf area lost per m^2^ of foliage) was analysed using a generalised linear mixed model with a beta error structure and model selection. As models with a beta error structure only allow values between 0 and 1, the 0 values in our study (n = 2) were converted into 0.0001 prior to analysis. We used herbivory as the response variable, while experimental treatment, elevation band and their interactions were predictor variables with tree species as a random effect. Again, elevation was modelled as a second-order polynomial. We used the glmmTMB package (Brooks, et al. 2017) within R 4.0.2 (Team 2020) to build our generalised linear mixed models.

### Body sizes

We modelled the mean body size of arthropods from each of the saplings using a similar model to that described above for arthropod density.

### Effect on groups of arthropods

We based this analysis on the unweighted natural log response ratios LRR (Curtis and Wang 1998, Hedges, et al. 1999) calculated from mean responses of individual groups of arthropods in the presence and absence of predators. LRR is an effect size measure that quantifies the results of experiments as the log-proportional change between the means of the treatment (in the absence of insectivorous predators, ⍰_I-_) and control group (in the presence of insectivorous predators, ⍰_I+_) and was thus calculated as (ln[⍰_I-_/ ⍰_I+_]) for each of the three treatments at each elevational study site and season. We then constructed linear models, selected the best one, and used the function *get_model_data* from package *metan* to obtain effect estimates (https://github.com/KatkaSam/GRAD_Exclosures).

Abundance, species richness, and biomass of insectivorous birds and bats were correlated (Pearson correlation) with the log response ratio (LRR, calculated from the raw data) of the vertebrate exclosure treatment at each elevation. The number of saplings with ants on them and abundance of ants on baits were correlated with log response ratio LRR of the ant exclosure treatment at each elevation. All data were normalized on the scale of 0-1 and averages across both seasons were taken as the data from the predator surveys did not exactly match the timing of the experimental predator exclusion.

## Results

### Change in arthropod densities

In general, the density of arthropods was higher and increased more in the wet season than in the dry season (Table S6). Based on the model predictions, the density of arthropods increased on average by 37% when the vertebrate predators were removed (Table S6). While the effect of vertebrates on arthropods was significant even when combined with the effect of ants (33% increase, Table S6), the exclusion of ants alone had no significant effect on arthropod density, which only increased by 12% based on the model predictions (Table S6). Ant removal was successful. There was a total of 538 and 884 ants found on control saplings (CON) and on saplings protected against vertebrates (VER) respectively, but only 13 and 30 on saplings protected against both types of predators (ALL) and ants only (ANT) respectively, across the whole experiment. Thus, ants represented 2.3% and 1.9% of all arthropods in CON and VER saplings respectively but less than 0.05% of arthropods on ANT and ALL saplings. Elevation and the interaction between elevation and treatment, and elevation and season, also had significant effects on the resulting density of arthropods (Figure 1, Table 1).

**Table 1.**
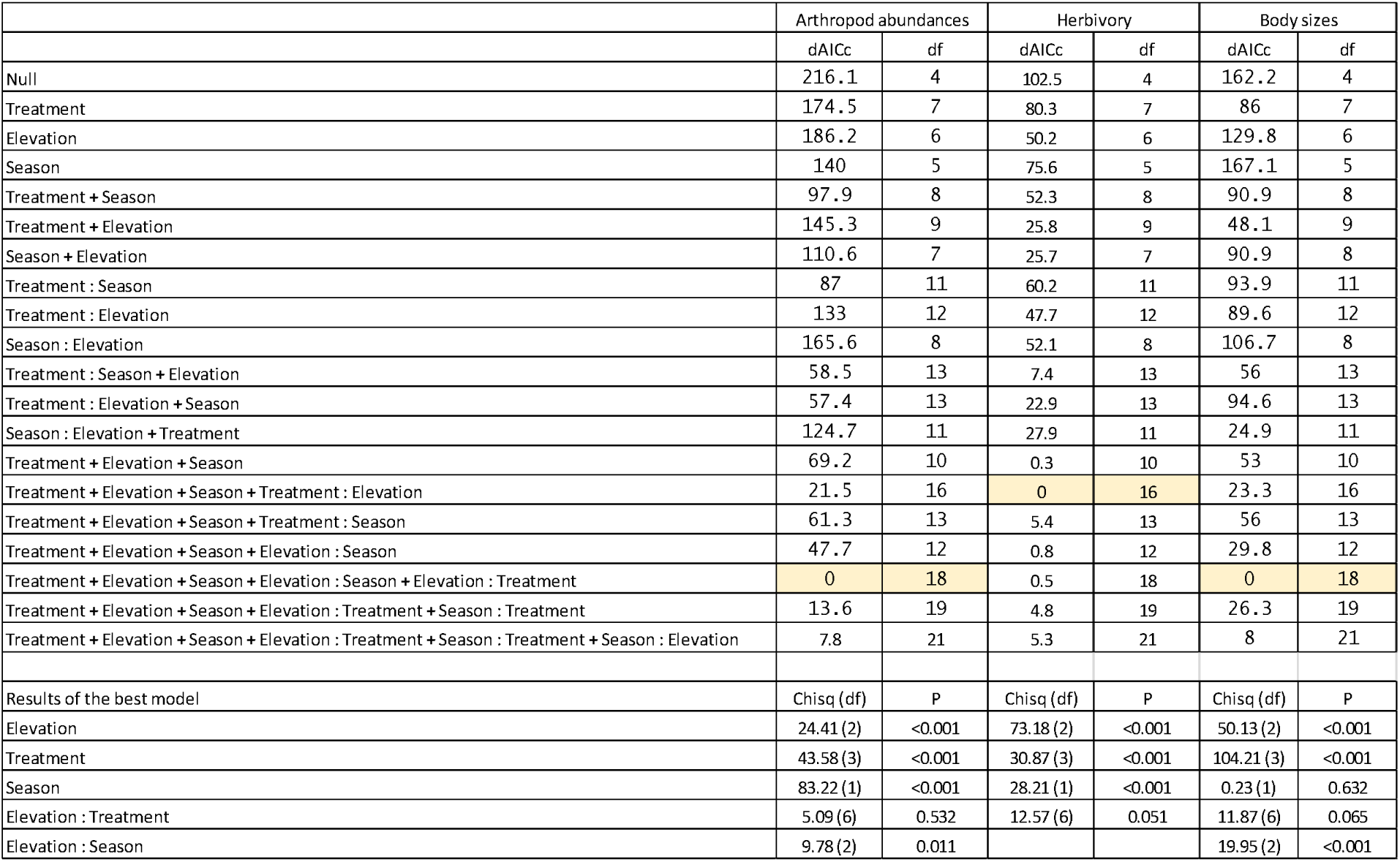
Corrected Akaike Information Criterion (AICc) of regression models examining abundances of all arthropods, herbivory damage, and mean body size of arthropods observed on saplings in the wet and dry season (season) and excluding various predators (treatment) along the elevational gradient of Mt. Wilhelm (elevation) in Papua New Guinea (a). Results of the analysis of variance of the most parsimonious models selected based on the AIC (b).

|  | Arthropod abundances |  | Herbivory |  | Body sizes |  |
| --- | --- | --- | --- | --- | --- | --- |
|  | dAICc | df | dAICc | df | dAICc | df |
| Null | 216.1 | 4 | 102.5 | 4 | 162.2 | 4 |
| Treatment | 174.5 | 7 | 80.3 | 7 | 86 | 7 |
| Elevation | 186.2 | 6 | 50.2 | 6 | 129.8 | 6 |
| Season | 140 | 5 | 75.6 | 5 | 167.1 | 5 |
| Treatment + Season | 97.9 | 8 | 52.3 | 8 | 90.9 | 8 |
| Treatment + Elevation | 145.3 | 9 | 25.8 | 9 | 48.1 | 9 |
| Season + Elevation | 110.6 | 7 | 25.7 | 7 | 90.9 | 8 |
| Treatment : Season | 87 | 11 | 60.2 | 11 | 93.9 | 11 |
| Treatment : Elevation | 133 | 12 | 47.7 | 12 | 89.6 | 12 |
| Season : Elevation | 165.6 | 8 | 52.1 | 8 | 106.7 | 8 |
| Treatment : Season + Elevation | 58.5 | 13 | 7.4 | 13 | 56 | 13 |
| Treatment : Elevation + Season | 57.4 | 13 | 22.9 | 13 | 94.6 | 13 |
| Season : Elevation + Treatment | 124.7 | 11 | 27.9 | 11 | 24.9 | 11 |
| Treatment + Elevation + Season | 69.2 | 10 | 0.3 | 10 | 53 | 10 |
| Treatment + Elevation + Season + Treatment : Elevation | 21.5 | 16 | 0 | 16 | 23.3 | 16 |
| Treatment + Elevation + Season + Treatment : Season | 61.3 | 13 | 5.4 | 13 | 56 | 13 |
| Treatment + Elevation + Season + Elevation : Season | 47.7 | 12 | 0.8 | 12 | 29.8 | 12 |
| Treatment + Elevation + Season + Elevation : Season + Elevation : Treatment | 0 | 18 | 0.5 | 18 | 0 | 18 |
| Treatment + Elevation + Season + Elevation : Treatment + Season : Treatment | 13.6 | 19 | 4.8 | 19 | 26.3 | 19 |
| Treatment + Elevation + Season + Elevation : Treatment + Season : Treatment + Season : Elevation | 7.8 | 21 | 5.3 | 21 | 8 | 21 |
| Results of the best model | Chisq (df) | P | Chisq (df) | P | Chisq (df) | P |
| Elevation | 24.41 (2) | <0.001 | 73.18 (2) | <0.001 | 50.13 (2) | <0.001 |
| Treatment | 43.58 (3) | <0.001 | 30.87 (3) | <0.001 | 104.21 (3) | <0.001 |
| Season | 83.22 (1) | <0.001 | 28.21 (1) | <0.001 | 0.23 (1) | 0.632 |
| Elevation : Treatment | 5.09 (6) | 0.532 | 12.57 (6) | 0.051 | 11.87 (6) | 0.065 |
| Elevation : Season | 9.78 (2) | 0.011 |  |  | 19.95 (2) | <0.001 |

**Figure 1.**
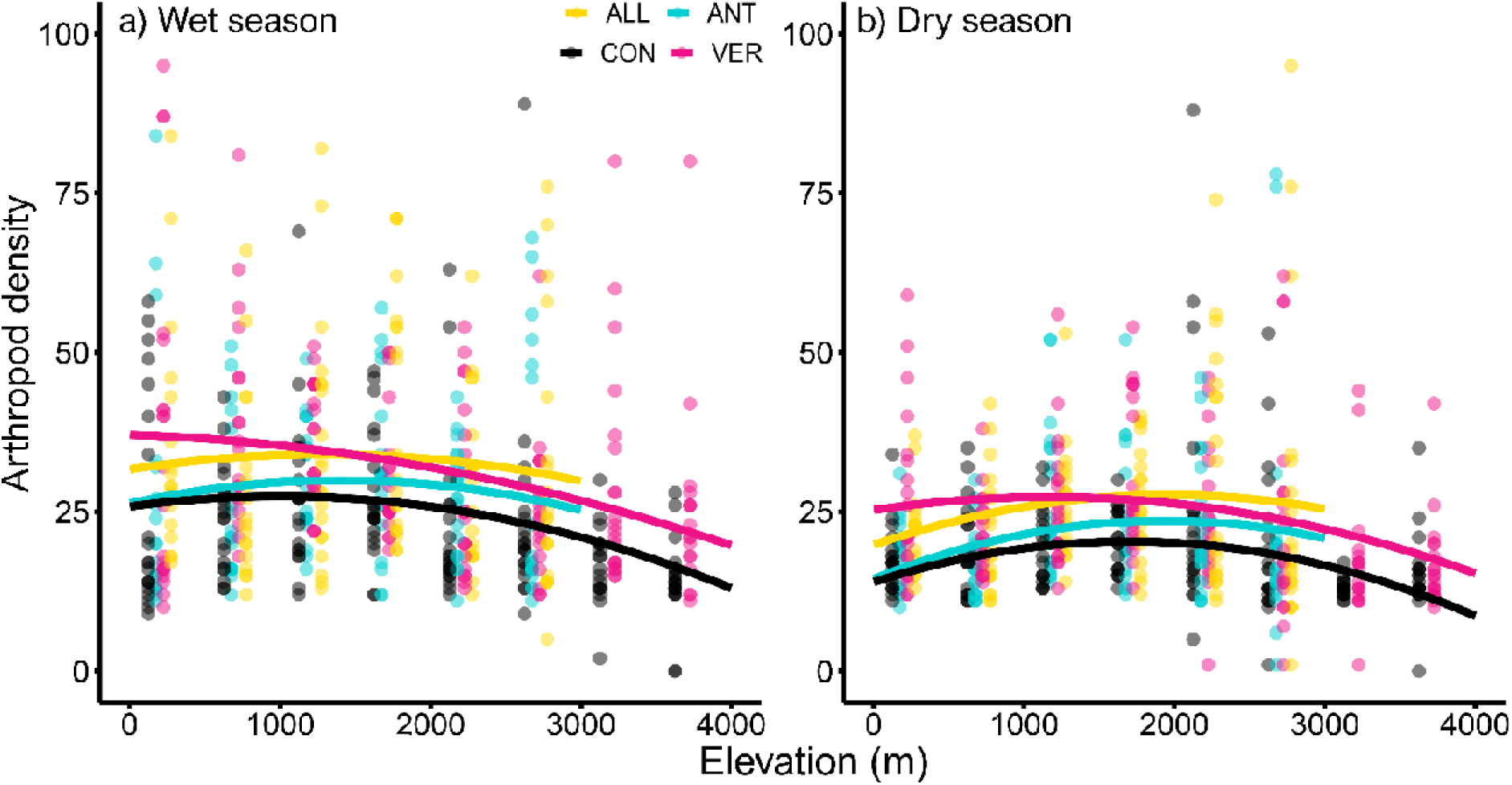
Effect of the predator exclusion (treatment) on the densities of all arthropods on saplings along the elevational gradient of Mt Wilhelm in the wet (a) and dry season (b). Each individual data point shows the resulting density of arthropods (per m^2^ of foliage) on a focal sapling. The curves show predicted values from the linear model which includes season, elevation and treatment, using the interaction between elevation and treatment as a fixed factor and plant species and identity of the sapling as a random effect. Treatments: CON = control saplings with free access of predators, ANT = saplings from which ants are excluded by tanglefoot glue, VER = saplings from which insectivorous vertebrates were excluded by cages, ALL = saplings from which both ants and insectivorous vertebrates were excluded.

### Change in herbivory

Predator exclusion, elevation and season had a significant additive effect on leaf area loss (Figure 2, Table 1). The natural mean standing herbivorous damage on control saplings at 200 m a.s.l. was 2.24 ± 1.7% in the wet season and 2.49 ± 1.7% in the dry season and decreased with increasing elevation [see Sam, et al. (2020) for more results]. Overall, exclusion of insectivorous vertebrates as well as the exclusion of both types of predators (ALL) led to a significant increase in leaf damage by 50 and 36% respectively (Table S7). However, the increase in leaf damage by ca. 9% in the ant exclosures was not significant (Table S7). Insects consumed significantly more leaf area in the wet than in the dry season (Figure 2). The increase in leaf damage seemed to be highest in the lowest and upper most elevations (Figure 2, Table S7).

**Figure 2.**
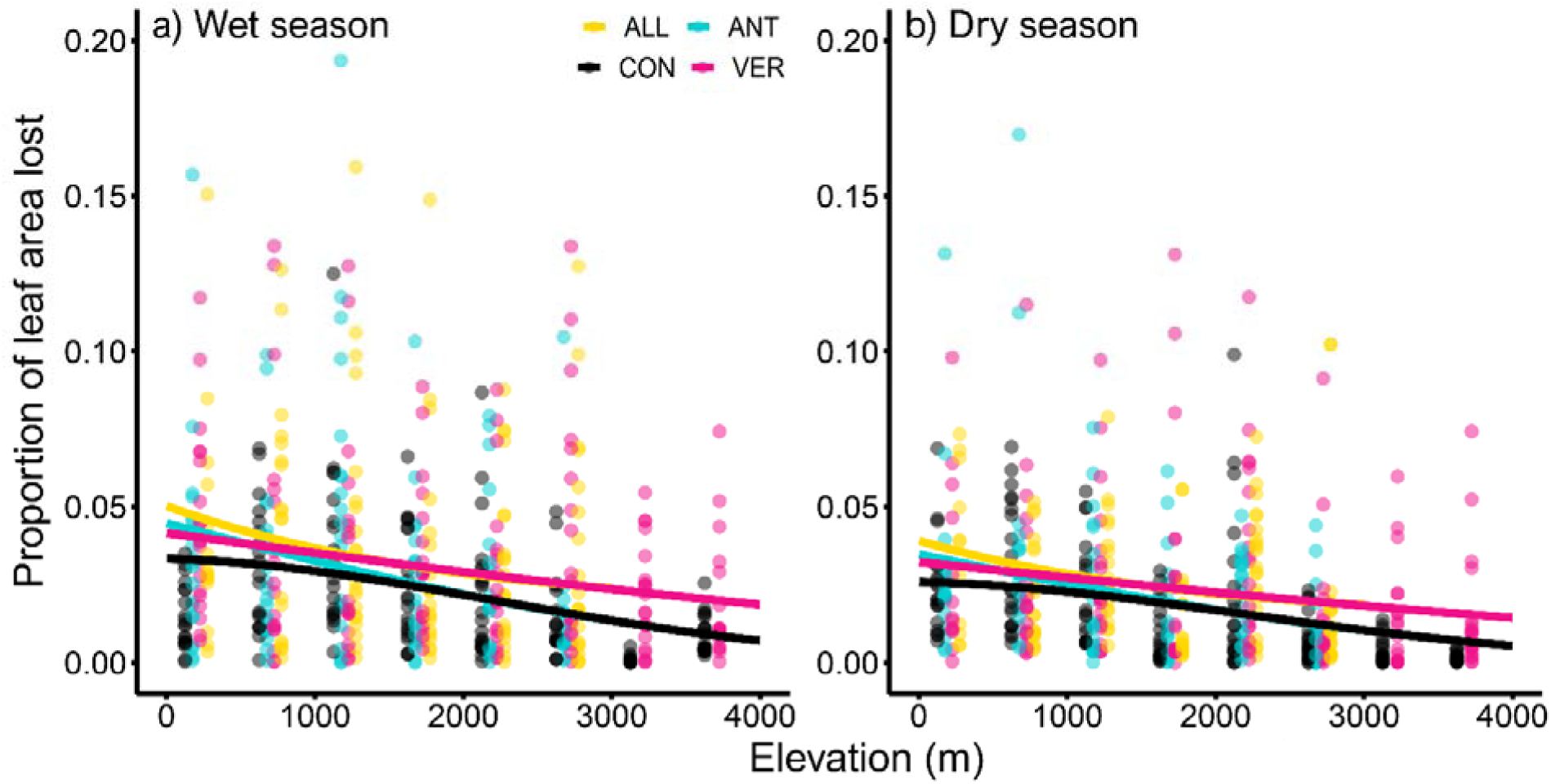
Effect of the predator exclusion (treatment) on leaf damage (m^2^ per sapling) on saplings along the elevational gradient of Mt Wilhelm in the wet (a) and dry season (b). Each individual data point shows the resulting herbivory damage of a focal sapling. The curves show predicted values from the glmmTMB models which include season, elevation and treatment as fixed factors and plant species and identity of the sapling as a random effect. Treatments: CON = control saplings with free access of predators, ANT = saplings from which ants are excluded by tanglefoot glue, VER = saplings from which insectivorous vertebrates were excluded by cages, ALL = saplings from which both ants and insectivorous vertebrates were excluded.

### Change in mean body size of arthropods

Treatment, elevation, season, and interaction between elevation and treatment, and elevation and season, had an impact on the mean body sizes of arthropods (Figure 3, Table 1). While model selection identified season as a significant factor due to its importance in model interactions, the effect of season alone was not significant (Table 1). This means that seasonal effect cannot be described properly because of its variation across elevations. The arthropods in caged treatments were significantly bigger than arthropods collected from control saplings and saplings from which ants were excluded (Figure 3, Table 1). The majority of arthropods collected during the experiment were very small, with larger arthropods (i.e,. greater than ca. 2.2 cm in length) found exclusively in the caged treatments (VER and ALL) to which vertebrate predators had no access.

**Figure 3.**
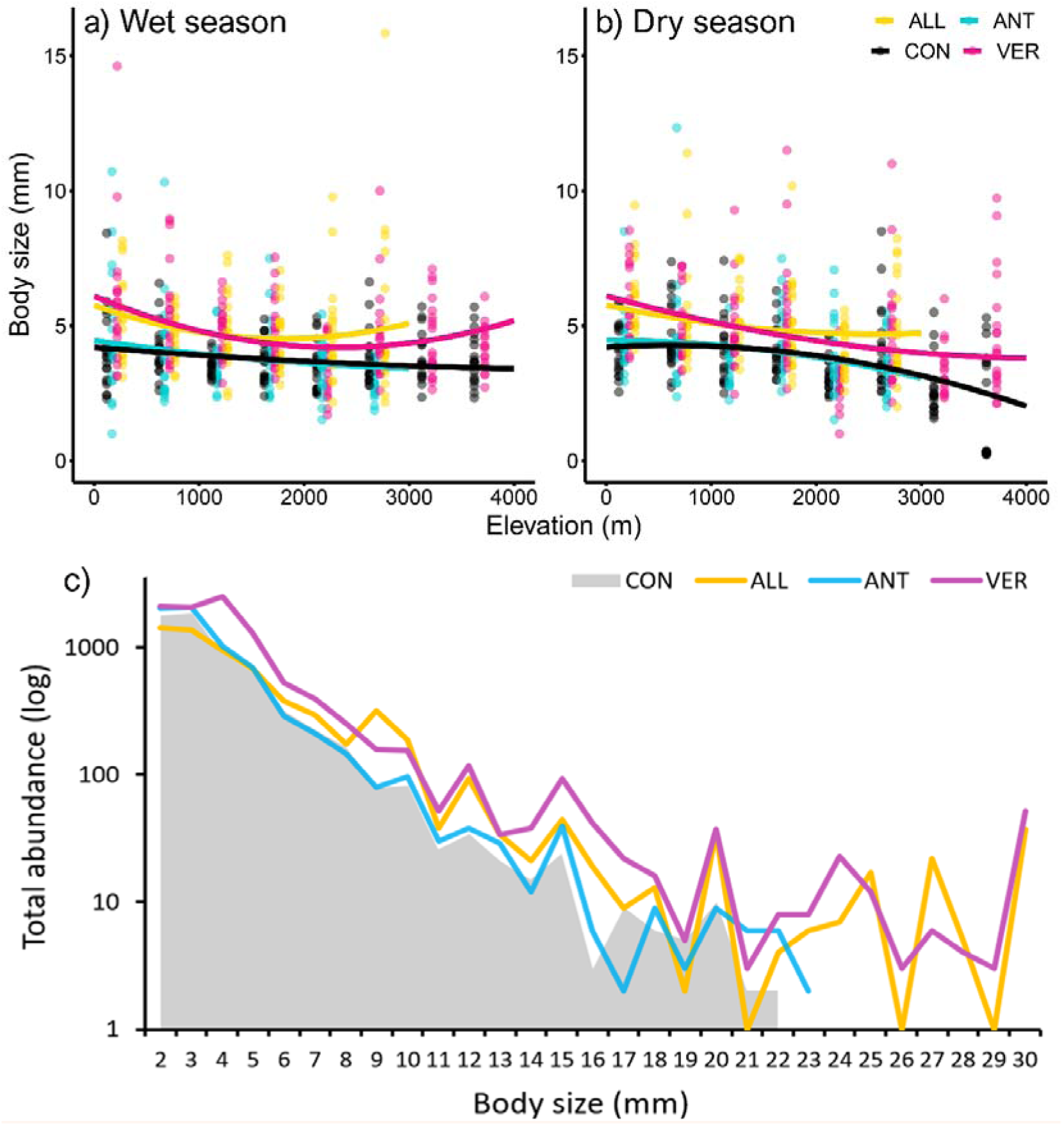
Effect of the predator exclusion on mean body size of the arthropods collected from saplings along the elevational gradient of Mt Wilhelm in the wet (a) and dry season (b). Each individual data point shows the resulting mean body size of the arthropod community on a given sapling. The curves show predicted values from the linear model which included season, elevation, and treatment as a fixed factor. See Table S6 for model results. The occurrence (log of the total abundance) of arthropods of a given body size in each of the four treatments is shown in (c). Treatments: CON = control saplings with free access of predators, ANT = saplings from which ants are excluded by tanglefoot glue, VER = saplings from which insectivorous vertebrates were excluded by cages, ALL = saplings from which both ants and insectivorous vertebrates were excluded.

### Changes in community composition

Overall, 12,177 arthropod individuals were identified as predatory arthropods, while only 4,085 were herbivores potentially responsible for leaf damage and 10,065 were arthropods which had no relationship to chewing herbivory (Table S3). The majority of predatory arthropods (8,221) were spiders. Absolute abundance as well as relative abundances of spiders were higher in treatments where ants were excluded (both ALL and ANT) but the change was not detectable in vertebrate exclosures (VER) (Figure 4). Across the whole study, Araneae, Diptera, Hemiptera, and Lepidoptera larvae increased their abundances significantly after the removal of ants. Araneae, Coleoptera, Lepidoptera larvae, Orthoptera and all other arthropods increased their densities (per m^2^ of foliage) on saplings from which the insectivorous predators were excluded. Densities of Hymenoptera (other than ants) and Hemiptera tended to decrease after vertebrate predators were excluded (Figure 4).

**Figure 4.**
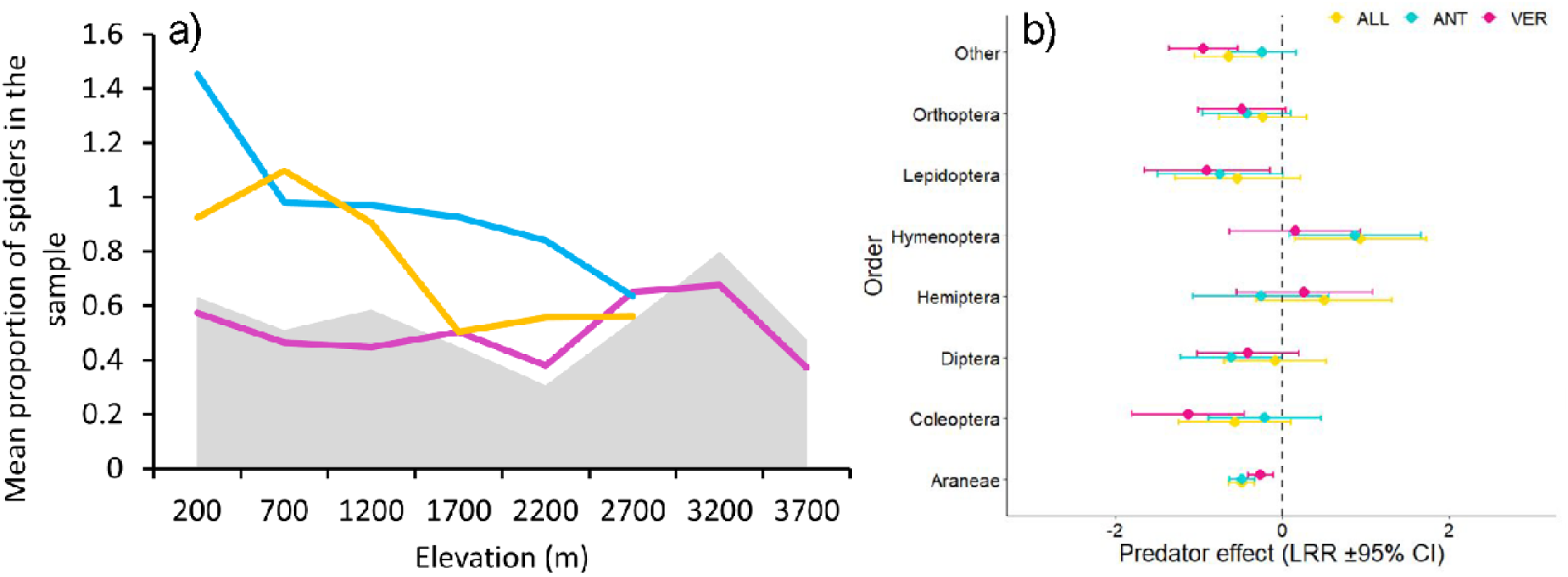
Mean proportion of spiders in control treatments (grey area), insectivorous vertebrate exclosures (VER - magenta line), ant exclosures (ANT - blue line) and both insectivorous vertebrates and ants exclosures (ALL-yellow line) across the elevational gradient (a). Caterpillar graph showing how individual orders of arthropods responded to the predator exclosure treatments. The X-axis shows mean effect sizes of natural log ratios (LRR = ln(exclosure/control)) with 95% CI of individual response variables. Effect size values above zero indicated that the absence of predators (treatment) was more harmful to plants than the control (presence of predators), as the abundances of arthropods in exclosures increased and caused potentially higher herbivory damage to plants (b). Treatments: ANT = saplings from which ants are excluded by tanglefoot glue, VER = saplings from which insectivorous vertebrates were excluded by cages, ALL = saplings from which both ants and insectivorous vertebrates were excluded.

### Predator communities and their relation to the observed cascading effect

Overall, we found only weak relationships between the communities of predators and the cascades they cause. In general, the abundances of predators correlated with the effect measured on abundances of prey (Figure 5), but not with the effect measured on herbivory damage (Figure S2). This may simply be the result of having just eight points in the correlation thus limiting the robustness of the analyses. Abundances of insectivorous birds, and thus those potentially responsible for attacks, peaked at mid elevations (1700 - 2700 m) and correlated positively and significantly (given the low power of the analysis and strong effect size) with the effect of vertebrate exclosure treatment on arthropod abundances (R = 0.71, P = 0.049, Figure 5a). Abundances of insectivorous bats decreased steeply from 200 to 1200 m a.s.l. and then peaked again at 2700m a.s.l. This pattern was again positively, but not significantly, correlated with the effect of vertebrate exclosure (R = 0.56, P = 0.15, Figure 5b). We detected more bats, and especially gleaning bats, in lowlands than at higher elevations [see also Sivault, et al. (2022) for more detailed results]. It seems that our results indicate that bats are at least partly responsible for predation at low elevations, while insectivorous birds are more important predators at the mid-elevations. Neither richness nor biomass of insectivorous birds or insectivorous bats correlated well with the effect of the exclosures measured on abundance of prey, and abundance of these insectivorous predators thus seemed to be a better correlate for predation pressure (R < 0.21, P > 0.78 in all cases). The number of trees on which ants occurred as well as ant abundances decreased with elevation. Despite the positive relationship, neither the number of trees with ants on them (R = 0.25, P = 0.56) nor abundance of the ants (R = 0.14, P = 0.74, Figure 5c) correlated significantly with the effect of the ant exclosures on abundances of arthropods.

**Figure 5.**
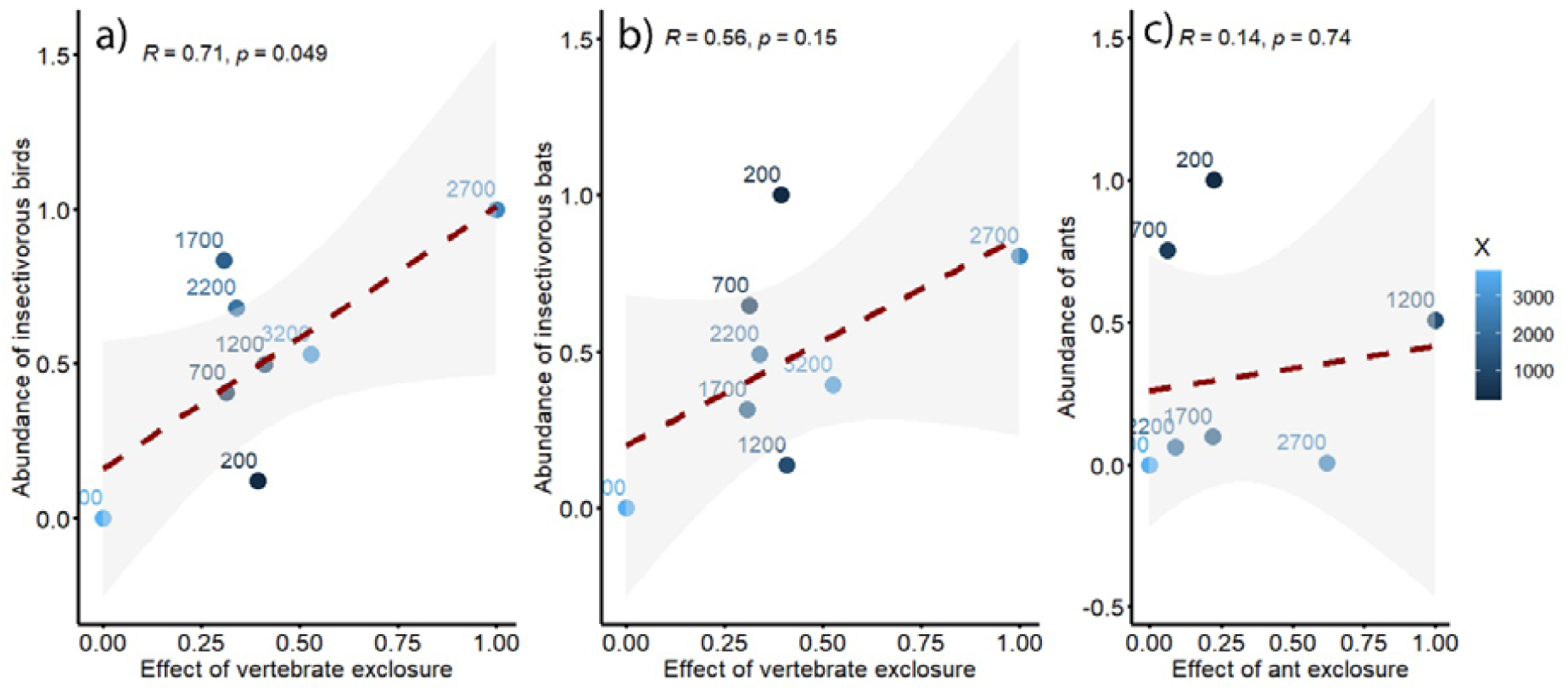
Correlations between the effect of vertebrate exclosures (a, b) and ant (c) exclosures (LRR calculated from raw data and normalized data) on abundances of arthropods and abundances (normalized) of insectivorous birds (a), insectivorous bats (b) and ants (c) potentially feeding on the experimental saplings.

## Discussion

Predator removal increases arthropod richness on plants, which is in line with our first hypothesis (*H1*). Overall, arthropod richness increased by ca. 37% after the removal of insectivorous vertebrates, which is in line with the estimates from two global reviews reporting an increase by ca. 39% (Mooney, et al. 2010) and 69 % (Sam, et al., 2022). The effect of ant removal was overall rather low, with arthropod abundance increasing by 12%, while the global effect of ant removal is estimated to be ca. 12% (Sam, et al., 2022) but also up to more than 100% (Rosumek, et al. 2009). However, it is important to note that Rosumek’s, et al. (2009) study analyses datasets comprising 70% myrmecophytic and only 30% non-myrmecophytic plants, while Sam’s, et al. (2022) analyses mostly data from non-myrmecophytic plants. Similarly, the plant species used in the current study have no or very weak associations with ants. It is important to note that the treatment itself almost completely removed ants from our experimental saplings, and ants represented ca. 2% of all arthropods in samples from our controls, thus impacting the effect of the treatment on the total density of the arthropods.

The overall effect of predators on lower trophic levels was however affected by season (*H1A*). Predator exclosures had a smaller effect on arthropod richness in the dry season than in the wet season. The seasonal differences are likely to be much higher in more seasonal habitats than those of the northern side of Papua New Guinea, which is known to be only weakly seasonal (Novotny and Basset 1998, Wright, et al. 1997). Birds also appear to follow the patterns observed here (Bell 1982). Observations made during our long-term surveys indicate that most breeding of insectivores occurs at the onset of rains, when the arthropods become more abundant in response to leaf flushing (Sam, et al. 2015, Sam, et al. 2017), which can lead to strong effects of predator exclosures.

Contrary to our expectations (*H1B*) the effect of insectivorous vertebrates on arthropod richness increased both towards the lowest (200 and 700m) and the highest elevations (above 2700m). Based on the productivity of the environment and previous observations on dummy prey predation rate by birds along the same gradient (Sam, et al. 2015), we expected that predator exclusion would have the highest effect between 700 and 1700 m a.s.l. where the abundances of insectivorous birds peaked. Our current results imply that the predation rate observed on dummy prey may not translate directly into the actual effect of predator exclusion, as there is an interplay between the natural abundances of arthropods and their predators. It is important to again note, that both groups of predators (birds and bats) were excluded in the current study. While the abundance of insectivorous bats was high at low and high elevations but low at mid-elevations, the abundance of insectivorous birds peaked at the mid-elevations. Moreover, bat communities between lowlands and highlands are different, with most of the gleaners found in lowlands. We can only speculate that bats are responsible, at least partly, for the predation at low elevations as our data are not suited to detect subtle elevational trends. We showed that predation risk on arthropods was correlated with the abundance of predatory birds (and less so for bats), supporting thus the hypothesis that predator density drives the effect of predation on lower trophic levels. These results are in line with some but not all earlier studies (Nordberg and Schwarzkopf 2019). Our data further indicates that the abundances of birds are a better correlate for the effect of predators than their richness or biomass.

Predator exclusion was affected not only arthropod densities but also their mean body sizes (*H2*). In general, the absence of insectivorous predators allowed larger arthropods to survive. This effect was detectable despite the fact that the mesh size (3×3 cm) of the cages was likely also preventing access of large arthropods. Arthropod mean body size increased primarily at low elevations where many large arthropods occur (Sam, et al. 2017, Horne, et al. 2018), but also at the highest elevations. In contrast to vertebrate insectivores, the exclusion of ants did not have any effect on the mean body size of arthropod communities. As such this contradicts results of an earlier study where ants preyed selectively upon small-bodied caterpillars, increasing thus mean caterpillar length by 6% (Singer, et al. 2017). However, we believe that this is because ants hunt not only individually for tiny prey, but also hunt collectively for large arthropods (Schmidt and Dejean 2018). This might lead to a balanced consumption of arthropods of all sizes.

Overall, the absence of predators allowed various groups of arthropods to increase their abundances (*H3*). First of all, insectivorous vertebrates did not affect ant density significantly, implying limited intraguild predation between these taxa in our study systems. The absolute numbers of ants increased on saplings protected against vertebrates, and ants represented very similar proportions of arthropods in samples from the control and vertebrate-protected saplings. Diet analyses of birds from the same gradient also showed that ants are not important prey for birds (Sam, et al. 2017). Other studies concur with this, where Coleoptera, Aranea, Lepidoptera larvae (Poulin, et al. 1994, Sam, et al. 2017) and many other smaller groups of arthropods (Bodawatta, et al. 2022, Poulin, et al. 1994) were found to be preferred prey of birds, but not ants (*H4*). This seems to be reflected in our current results, as these groups of arthropods increased their abundances significantly after the insectivore predators were removed, while other groups (e.g., Hymenoptera, Diptera, Hemiptera) did not respond strongly to the absence of predators. In the ant exclosures, Hemiptera, Lepidoptera, and Aranea increased their abundances significantly, but spiders seemed to be the most affected. This is in line with results of earlier studies which showed that spiders increased their abundances by more than 100% (Rosumek, et al. 2009) and by 84% (Sam, et al. 2022) in the absence of ants. Even though our results can’t resolve whether the spiders are typical prey of ants or their competitors, we believe that competition might be at least partially responsible for the patterns observed as spider abundance was higher in the absence of ants than in the absence of vertebrates. Results of existing studies are vague, and despite most of the studies typically show ants to feed on eggs and insect larvae primarily (Gathalkar and Sen 2018), there is some evidence of them hunting spiders also (Risch and Carroll 1982).

In contrast to previous reports, the standing herbivory on control saplings along Mt Wilhelm was 5 to 10 times lower than the global estimates and we discuss this matter and potential reasons in an earlier study (Sam, et al. 2020). The change in herbivory following the removal of predators was also surprising (*H5*). Herbivory damage increased significantly both after removal of ants and vertebrate predators but varied in time and space. Herbivory across the whole gradient in the wet season increased by 112, 78 and 78% after the removal of all predators, vertebrate predators, and ants respectively. In contrast, the dry season saw increases of 58, 52 and 42% respectively. All of these increases are higher than the previous global estimates which predicted increases in leaf herbivory of ca. 40 - 47% in the absence of vertebrate predators (Mooney, et al. 2010, Sam, et al., 2022). The effect of ants on herbivory at low elevations was actually stronger than their effect on the density of herbivore arthropods, where ants increased herbivory by ca. 130% and herbivorous arthropods only by 3%. Similarly, Rosumek et al. (2009) showed that ant removal increased herbivory by more than 95%, whereas the abundances of herbivorous arthropods increased by ca. 60% in the tropics.

Season had an effect on both the densities of arthropods and herbivory. The removal of vertebrate predators in the wet season led to increases in herbivory of ca. 100-230%, where peaks towards the lowest and higher elevations created a distinct U-shaped pattern. We saw the highest increase in herbivory during the dry season following the removal of vertebrates above 2200m which ranged between 160% and 555%. This pattern suggests that insectivorous predators provide more effective services to plants at the highest and lowest than at middle elevations. However, this is likely heavily dependent on plant defences and bottom-up control of the trophic cascades.

According to one hypothesis, plants are expected to invest progressively less into chemical defences at higher altitudes as arthropod richness is generally lower there (Garibaldi, et al. 2011, Pellissier, et al. 2014). Our results did not support this however, as we did not observe a steep decrease in arthropod densities along our elevational gradient. A second hypothesis states that replacing biomass at higher elevations comes at a higher cost due to the harsh, low nutrients environments of higher elevations. These conditions favour higher investment into defences at the expense of growth (Defossez, et al. 2018, Salgado, et al. 2016). Plants may also defend their biomass directly and indirectly through insectivorous predators (Marquis and Whelan 1996). We might speculate that plants do not invest much into direct defences at higher elevations as arthropods are relatively rare there, and insectivorous birds more abundant, but instead rely on indirect defences which work in response to herbivory attacks. While this would explain the pattern we detected, it appears unlikely given our understanding of plant defences. In an earlier study from the same elevational gradient using several of the plant species studied here (Sam, et al. 2020), showed that there is no universal elevational increase in plant defences (i.e., levels of alkaloids and flavonoids), and several Ficus species from mid elevations were also well defended (i.e., high protein precipitation capacity). If we wish to better understand elevational trends in plant defences then it is necessary to perform analyses based on multiple traits of direct and indirect defences and to link these to datasets on arthropods herbivores.

In conclusion, we show that vertebrate insectivores suppress arthropods effectively, yet their impact differs among various groups of arthropods. The pressure of vertebrate insectivores is robust across the whole elevational gradient and translates to a significantly higher plant herbivory in the absence of predators. Our results imply that the relative contribution of birds and bats differs along the elevational gradient. In contrast, ants do not suppress arthropods significantly, yet the trophic cascades caused by them do translate into increased herbivory at the most productive sites of the elevational gradient, where the abundances of ants are also the highest. Insectivorous vertebrates, but not ants, had a stronger negative effect on large arthropods which can further negatively impact herbivory rates. While generalities emerge, we suggest that future experiments need to consider intraguild predation and mesopredators and plant defences, and that these should be accompanied by more detailed predator surveys as all these factors may significantly affect the strength of trophic cascades.

## Data availability statement

All data and scripts are available at GitHub https://github.com/KatkaSam/GRAD_Exclosures

### Bioscatch

Katerina Sam is a head of Laboratory of Multitrophic interactions and the Department of Ecology at the Entomology Institute, where she studies relationships between insectivorous predators, insect and plants. This work represents her earlier postdoctoral experiment, which further grew into a European Research Council funded project. She and her colleagues study various aspects of ecology and biodiversity of Mt. Wilhelm from 2008.

## Acknowledgements

We are thankful to villagers from Kausi, Numba, Degenumbu, Sinopass, Bruno Sawmill and Kegesugl for allowing work on their land and assistance in the field. The work wouldn’t be possible without the logistical support and help of The New Guinea Binatang Research Centre. We are thankful for various support of Prof. Vojtech Novotny, Prof. Nigel Stork and Prof. Roger Kitching for support during the field and laboratory work, which was carried during the maternal leave of KS. The work was supported by Czech Science Foundation Grant 18-23794Y and European Research Council Starting Grant BABE 805189 (PI: K. Sam).

## Author contributions

KS conceived the ideas, secured funding and conducted the experiment with help of BK. PA and ES conducted the survey of insectivorous predators, LRJ assisted with statistical analyses to KS. All authors contributed to writing and edited the first draft written by KS.

## Notes

### Competing Interest Statement

The authors have declared no competing interest.

### Summary of Updates

Only English and grammar was improved by native speaker in the verison, and several sentences restructured for clarity.

