## Supplement for "Vertebrates but not ants protect rainforest trees from herbivorous insects along an elevational gradient in Papua New Guinea"

Supplementary material

**Figure S1.** Map of the Mt. Wilhelm elevational gradient showing all eight elevational study sites. Insert shows location of the elevational range within Papua New Guinea. Note that data from six lower elevational study sites were analysed separately from the full elevational gradient due to different plant species composition.

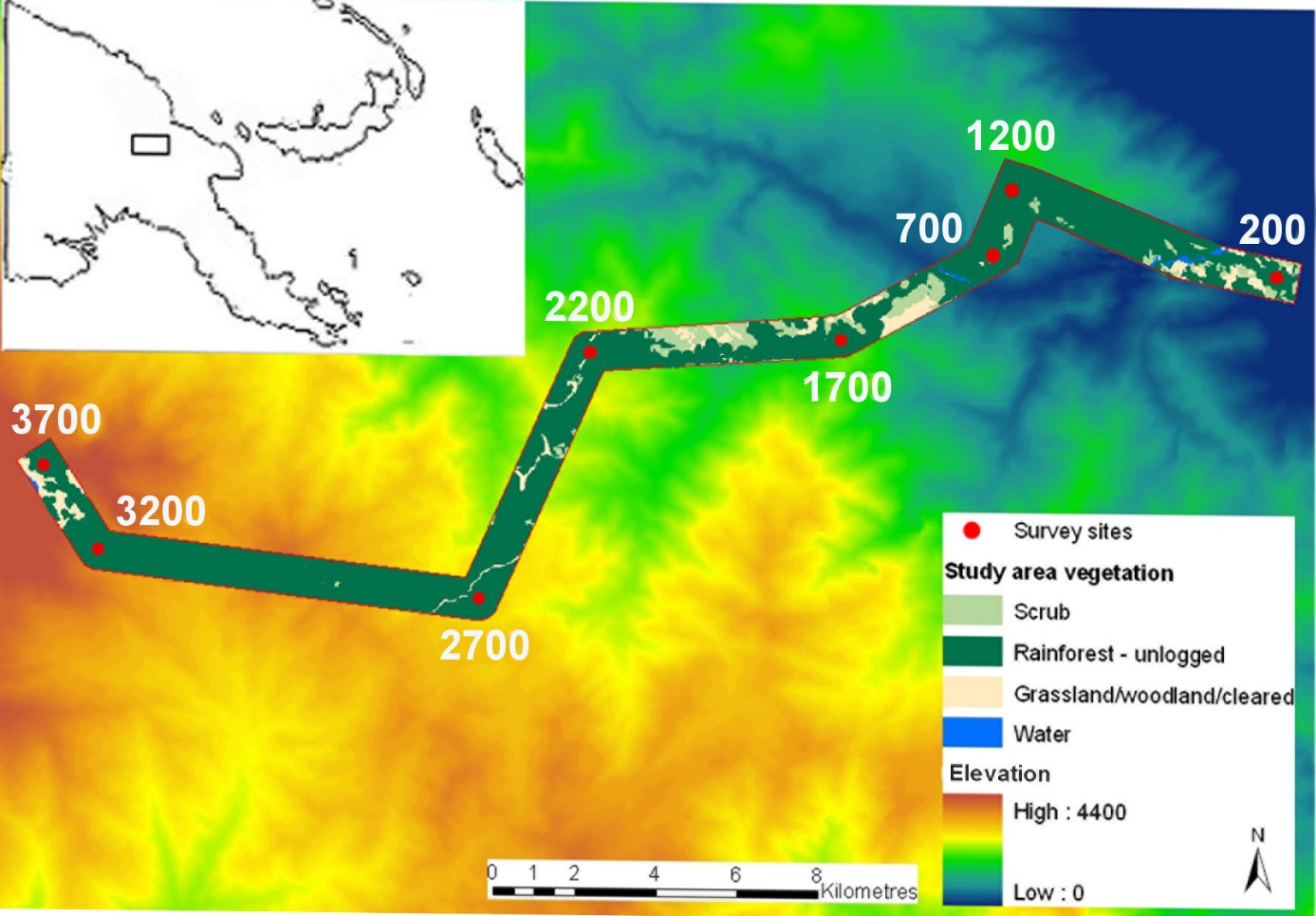

**Table S1.** Plant species, treatments, and the number of samples (total = 1,115) collected from each treatment from them at each elevational study site. ALL = vertebrate predators and ants excluded, ANT = ants excluded, VER = vertebrates excluded, CON = control saplings without exclusion.

| **Plant species Treatment** | | **200** | | **700** | | **1200** | | **1700** | | **2200** | | **2700** | | **3200** | | **3700** |
| --- | --- | --- | --- | --- | --- | --- | --- | --- | --- | --- | --- | --- | --- | --- | --- | --- |
| *Ficus arfakensis* | ALL | 8 | 4 | |  | | 14 | |  | |  | |  | |  | |
|  | ANT | 8 | 6 | |  | | 12 | |  | |  | |  | |  | |
|  | CON | 8 | 4 | |  | | 12 | |  | |  | |  | |  | |
|  | VER | 10 | 6 | |  | | 18 | |  | |  | |  | |  | |
| *Ficus badiopurpurea* | ALL |  | 8 | |  | | 6 | |  | |  | |  | |  | |
|  | ANT |  | 8 | |  | | 4 | |  | |  | |  | |  | |
|  | CON |  | 4 | |  | | 8 | |  | |  | |  | |  | |
|  | VER |  | 7 | |  | | 2 | |  | |  | |  | |  | |
| *Ficus congesta* | ALL | 6 |  | |  | |  | |  | |  | |  | |  | |
|  | ANT | 12 |  | |  | |  | |  | |  | |  | |  | |
|  | CON | 10 |  | |  | |  | |  | |  | |  | |  | |
|  | VER | 12 |  | |  | |  | |  | |  | |  | |  | |
| *Ficus conocephalifolia* | ALL | 8 | 4 | |  | |  | |  | |  | |  | |  | |
|  | ANT | 4 | 4 | |  | |  | |  | |  | |  | |  | |
|  | CON | 4 | 8 | |  | |  | |  | |  | |  | |  | |
|  | VER | 4 | 4 | |  | |  | |  | |  | |  | |  | |
| *Ficus endochaete* | ALL |  |  | | 24 | |  | | 8 | | 6 | |  | |  | |
|  | ANT |  |  | | 28 | |  | | 2 | | 4 | |  | |  | |
|  | CON |  |  | | 36 | |  | | 2 | | 4 | |  | |  | |
|  | VER |  |  | | 28 | |  | | 6 | | 8 | |  | |  | |
| *Ficus hahliana** | ALL | 12 | 3 | | 2 | |  | | 2 | | 12 | |  | |  | |
|  | ANT | 8 | 10 | | 6 | |  | | 6 | | 12 | |  | |  | |
|  | CON | 14 | 6 | | 2 | |  | | 4 | | 12 | |  | |  | |
|  | VER | 8 | 2 | | 2 | |  | | 6 | | 6 | |  | |  | |
| *Ficus hispidioides* | ALL | 4 |  | |  | |  | |  | |  | |  | |  | |
|  | ANT | 8 |  | |  | |  | |  | |  | |  | |  | |
|  | CON | 4 |  | |  | |  | |  | |  | |  | |  | |
|  | VER | 6 |  | |  | |  | |  | |  | |  | |  | |
| *Ficus hombroniana*^ǂ^ | ALL |  | 4 | |  | | 4 | | 10 | |  | |  | |  | |
|  | ANT |  | 4 | |  | | 2 | | 12 | |  | |  | |  | |
|  | CON |  | 2 | |  | | 8 | | 4 | |  | |  | |  | |
|  | VER |  | 2 | |  | | 4 | | 4 | |  | |  | |  | |
| *Ficus iodotricha* | ALL |  |  | |  | | 4 | | 4 | | 4 | |  | |  | |
|  | ANT |  |  | |  | | 10 | | 2 | | 9 | |  | |  | |
|  | CON |  |  | |  | | 4 | | 6 | | 2 | |  | |  | |
|  | VER |  |  | |  | | 6 | | 12 | | 6 | |  | |  | |
| *Ficus mollior* | ALL |  |  | |  | | 6 | |  | |  | |  | |  | |
|  | ANT |  |  | |  | | 4 | |  | |  | |  | |  | |
|  | CON |  |  | |  | | 6 | |  | |  | |  | |  | |
|  | VER |  |  | |  | | 2 | |  | |  | |  | |  | |
| *Ficus saccata* | ALL |  |  | |  | |  | | 6 | | 18 | |  | |  | |
|  | ANT |  |  | |  | |  | | 2 | | 14 | |  | |  | |
|  | CON |  |  | |  | |  | | 11 | | 21 | |  | |  | |
|  | VER |  |  | |  | |  | | 8 | | 20 | |  | |  | |
| *Ficus subcuneata* | ALL |  | 14 | |  | |  | |  | |  | |  | |  | |
|  | ANT |  | 2 | |  | |  | |  | |  | |  | |  | |
|  | CON |  | 8 | |  | |  | |  | |  | |  | |  | |
|  | VER |  | 12 | |  | |  | |  | |  | |  | |  | |
| *Ficus trichocerasa^●^* | ALL |  | 2 | | 14 | | 6 | |  | |  | |  | |  | |
|  | ANT |  | 6 | | 7 | | 7 | |  | |  | |  | |  | |
|  | CON |  | 8 | | 2 | | 2 | |  | |  | |  | |  | |
|  | VER |  | 6 | | 10 | | 8 | |  | |  | |  | |  | |
| *Ficus wassa*^#^ | ALL |  |  | |  | |  | | 10 | |  | |  | |  | |
|  | ANT |  |  | |  | |  | | 16 | |  | |  | |  | |
|  | CON |  |  | |  | |  | | 14 | |  | |  | |  | |
|  | VER |  |  | |  | |  | | 4 | |  | |  | |  | |
| *Pittosporum berberidoides* | CON |  |  | |  | |  | |  | |  | | 10 | | 16 | |
|  | VER |  |  | |  | |  | |  | |  | | 10 | | 16 | |
| *Myrsine womersleyi* | CON |  |  | |  | |  | |  | |  | |  | | 12 | |
|  | VER |  |  | |  | |  | |  | |  | |  | | 10 | |
| *Myrsine papuana* | CON |  |  | |  | |  | |  | |  | | 18 | | 6 | |
|  | VER |  |  | |  | |  | |  | |  | | 10 | | 10 | |
| *Macaranga melanosticta* | CON |  |  | |  | |  | |  | |  | | 12 | | 6 | |
|  | VER |  |  | |  | |  | |  | |  | | 20 | | 4 | |

Notes to Table S1: ^*^*F. hahliana* is confirmed as a good species from 200-1200m of our elevational gradient. After this (1700m-2700m) a close relative/sister species occurs. However, this potential split was discussed only recently based on molecular differences (Segar et al. 2016). We were not able to distinguish the two species in the time of our experiment. ^ǂ^ *F. hombroniana* is found between 200-1200m. There are a few individuals at 1,700m but most individuals classified as *F. hombroniana* here are probably (and at 2200m) the closely related *F. ihuensis. ^●^ F. trichocerasa* has two sub-species along the elevational gradient. *F. trichocerasa* subsp. *trichocerasa* occurs between 200-1700m and *F. trichocerasa* subsp. *pleioclada* occurs between 1700m and 2200m. They co-occur at 1700m, and both subspecies were included in our study as they are difficult to distinguish in the field at 1700m. ^#^ *F. wassa* comprises several varieties along the elevational gradient of Mt. Wilhelm (Berg and Corner 2005). The varieties included in our study were *F. wassa* var. nubigena which occurs along the gradient from 1300 to 3000 m and *F. wassa* var. wassa which occurs along the whole gradient.

**Table S2.** Number of saplings set for the treatments at individual elevational study sites; the first day when the treatments were set (T_0_), when the effect was surveyed for the first time and second time (S1, S2). We selected total of 560 saplings for the experiment, not all sapling however survived until the end of the experiment which leads to discrepancies between Table S2 and Table S1 (1,115 samples collected instead of 1,120 predicted). Treatments: CON – control trees ANT - only ants excluded, VER – only vertebrates excluded, ALL – vertebrates and ants excluded. Setting and survey of the exclosures usually took 2-4 days and was conducted by two teams working simultaneously or together according to safety, terrain, and safety needs.

| Elevation | CON | ANT | VER | VER+ANT |
| --- | --- | --- | --- | --- |
| 200 | 20 | 20 | 20 | 20 |
|  | T_0_:1May14 S1:1Nov14 S2:15Apr15 | | | |
| 700 | 20 | 20 | 20 | 20 |
|  | T_0_:29Apr14 S1:27Oct14 S2:7Apr15 | | | |
| 1200 | 20 | 20 | 20 | 20 |
|  | T_0_:25Apr14 S1:21Oct14 S2:30Mar15 | | | |
| 1700 | 20 | 20 | 20 | 20 |
|  | T_0_:27Apr14 S1:25Oct14 S2:8May15 | | | |
| 2200 | 20 | 20 | 20 | 20 |
|  | T_0_:14Apr14 S1:6Oct14 S2:16Mar15 | | | |
| 2700 | 20 | 20 | 20 | 20 |
|  | T_0_:11Apr14 S1:8Oct14 S2:19Mar15 | | | |
| 3200 | 20 | NA | 20 | NA |
|  | T_0_: 4Apr14 S1:31Aug14 S2: 16Feb15 | | | |
| 3700 | 20 | NA | 20 | NA |
|  | T_0_: 2Apr14 S1:31Aug14 S2:9Feb15 | | | |

**Table S3. List of the insect groups identified within each feeding guild, their abundances on individual treatments in wet and dry season. Predators =** arthropods able to kill other arthropods,

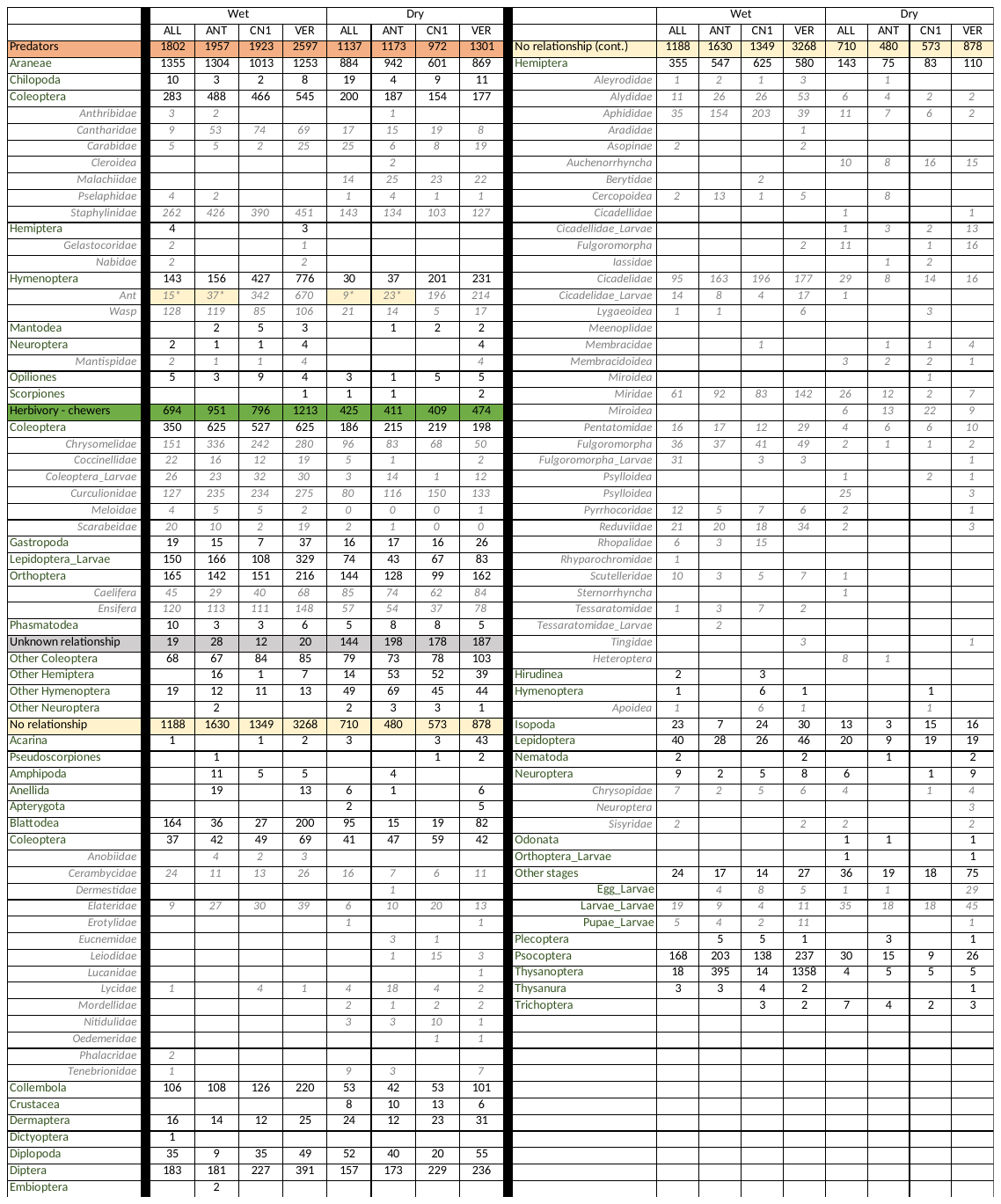

**Additional methods – survey of predators**

**Bird survey**

Bird communities here have been repeatedly surveyed by point counts as part of long-term monitoring efforts, with this methodology and data published in several studies (Marki, et al. 2016, Sam and Koane 2014, Sam, et al. 2019). In short, point counts at each elevational site were carried out at 16 points regularly spaced along a 2,350 m transect (successive points were 150 ± 5 m apart to avoid overlap). All birds seen or heard within a fixed radial distance of 0 - 50 m were recorded. Each count lasted 15 min so that all 16 points were surveyed before 11:00 (i.e., such survey on all 16 points represents one replication in time). All points and all study sites were surveyed equally during two independent surveys.

Here we used data from two surveys conducted prior to and during the exclosure experiment. The first survey was conducted between 15^th^ September and 15^th^ October 2013, i.e., 3 first consecutive days from a larger study by Sam, et al. (2019). The second survey was conducted between 1^st^ October 2015 and 2^nd^ December 2015 (again 3 consecutive days per site). Our analyses here use abundances and species richness of 217 forest bird species recorded during the standardized point-counts, which represents 76.5% of the avifauna known from the region (Marki, et al. 2016). We included the earlier data from 2013 to calculate mean abundances of insectivorous birds more robustly, as a 3-day long survey alone might not be sufficiently describing these tropical bird communities. Birds (Table S4) were partitioned into five trophic guilds: insectivores, frugivores, frugivore-insectivores, insectivore-nectarivores and nectarivores, based on dietary information extracted from the literature (Hoyo, et al. 1992-2011, Pratt and Beehler 2015, Sam, et al. 2017). We also extracted the body mass of each species and used this to calculate total bird biomass at each elevation.

**Bat survey**

Bat communities were surveyed in the understory during two expeditions conducted in wet (February – March 2015) and dry seasons (June – July 2015). This method and data were published in Sivault, et al. (2022). In short, we used an ultrasonic bat call detector coupled with a recorder to detect echolocating bat species. We recorded bats at five points (i.e., 15 minutes per point) separated by 200 meters at each elevation, in line with the bird transect methods described above. Surveys were conducted for four days per site after sunset (6 pm) but were only feasible for two days at 3,200 and 3,700 m due to logistical problems. Recordings were analysed with Adobe Audition version 22.0. We distinguished individual echolocation call types within the recordings and later identified these based on information from other surveys (Sivault, et al. 2022). All bat species identified (Table S5) were perceived as strict insectivores based on the literature (Bonaccorso 1998, Flannery 1995). Despite a low sampling effort, we estimated bat activity as a proxy of relative abundance. We retained one bat pass per five-second interval, which is the mean duration of all bat species passes as indicated by (Kerbiriou, et al. 2019). Finally, we calculated biomass of bats at each elevation using the average maximum body mass of males and females of each bat species found in the literature (Bonaccorso 1998, Flannery 1995). Any missing body mass information was supplemented with the body mass of the closest species(i.e., similar head-body size and/or forearm length).

**Ant survey**

The ant communities at each of the six study sites were sampled by hand collection. The trunk of each sapling was inspected for ants at breast-height for 10 minutes, and also by the tuna bait method in May-June 2014. Baits were filled with commercial canned tuna in oil, which is a standard method in studies of foraging ant communities. One teaspoon of tuna was placed as a bait under a strip of gauze at breast height on each of the saplings. Baits were inspected three hours following their exposure. The abundance of ants was counted on each bait. The combination of both methods was used to account for the fact that not all ant species are attracted to baits (Véle, et al. 2009). We correlated ant abundance data collected by tuna baits from understory saplings and hand collected data from the trunks. The same survey was conducted at 3200 and 3700 m above sea level and detected no ants, which is in line with results from previous studies of ant communities along the same gradient (Colwell, et al. 2016, Moses, et al. 2021, Sam, et al. 2015).

**Table S4.** List of recorded bird species, their feeding specialization, occurrence in forest strata (in %), body weight and abundance at each of the surveyed study sites.

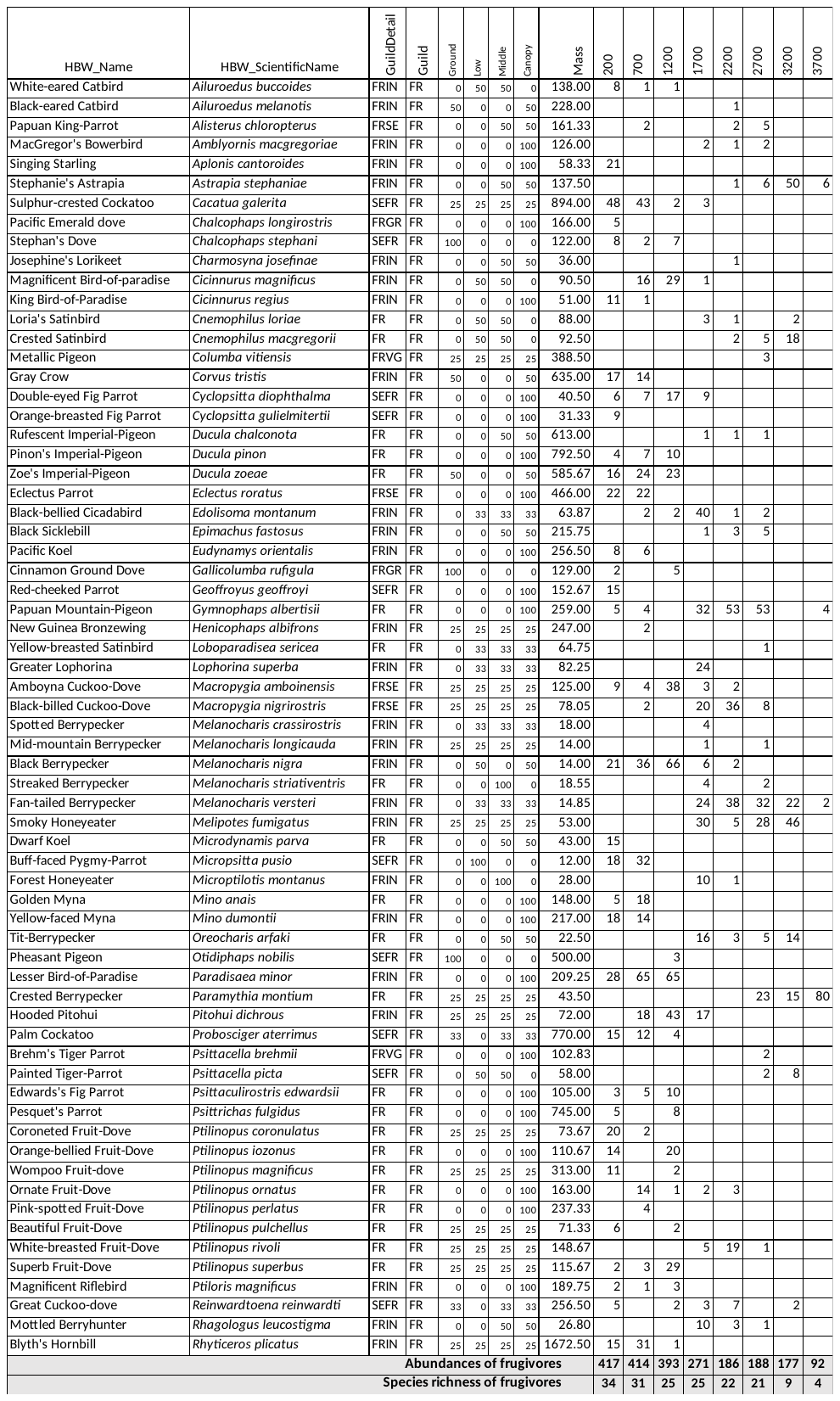

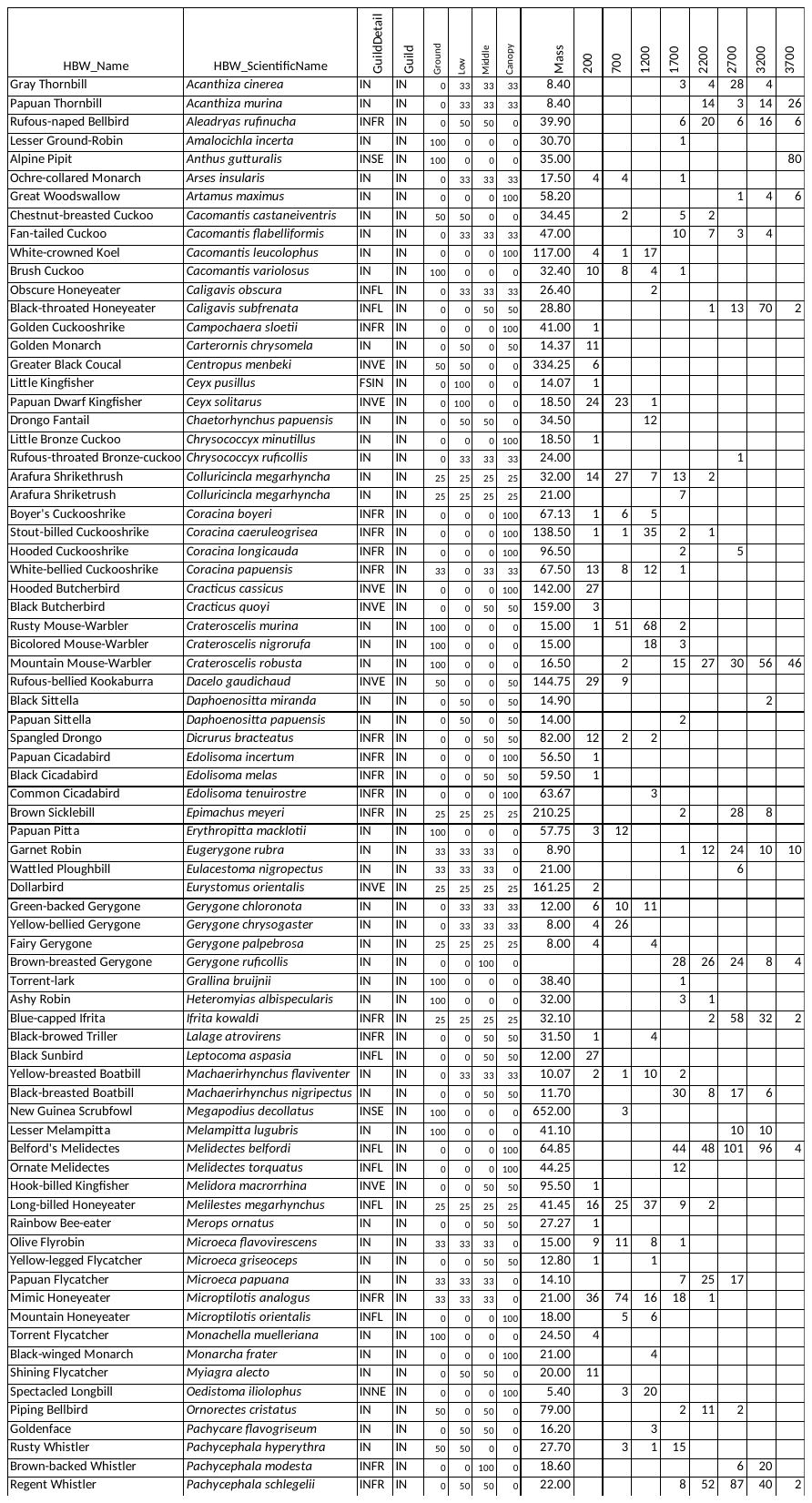

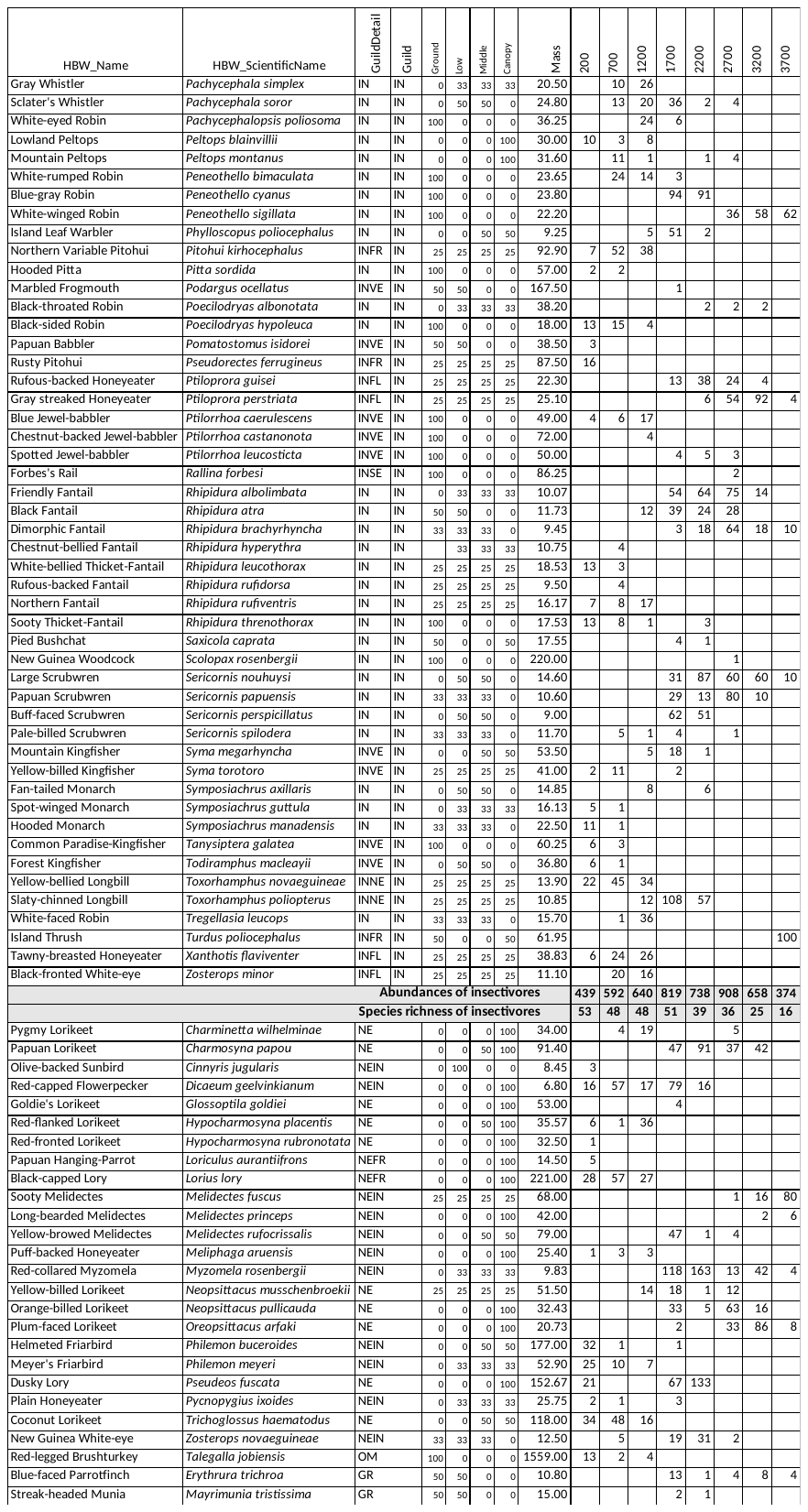

**Table S5. List of recorded bats and their feeding specializations.** Bat species acoustically recorded along the Mt. Wilhelm gradient. Numbers represents the relative abundance calculated from the recordings. Feeding guilds and foraging habits come from the literature (Flannery 1995, Bonaccorso 1998, Zachos et al. 2020).

| **Species** | **Call type** | **200** | **700** | **1,200** | **1,700** | **2,200** | **2,700** | **3,200** | **3,700** | **Food** | **Foraging** |
| --- | --- | --- | --- | --- | --- | --- | --- | --- | --- | --- | --- |
| **HIPPOSIDERIDAE** |  |  |  |  |  |  |  |  |  |  |  |
| *Hipposideros wollastoni* | 84 mCF |  |  |  |  | 22 |  |  |  | Likely insects | NA |
| *Hipposideros cervinus* | 140 sCF | 2 |  |  |  |  |  |  |  | Beetles, moths and other insects | Aerial and gleaning |
| **EMBALLONURIDAE** |  |  |  |  |  |  |  |  |  |  |  |
| *Emballonura beccarii /*  *Mosia nigrescens* | 62 i.fFM.d | 12 | 16 | 10 | 1 |  |  |  |  | Beetles, wingless ants | Aerial and gleaning |
| *Emballonura beccarii /*  *Mosia nigrescens* | 72 i.fFM.d | 12 |  |  |  |  |  |  |  |  |  |
| **VESPERTILIONIDAE** |  |  |  |  |  |  |  |  |  |  |  |
| *Pipistrellus papuanus* | 48 st.cFM | 10 |  |  |  |  |  |  |  | Aerial insects | Hawker |
| *Nyctophilus microtis* | 45-50 bFM | 5 | 4 | 1 |  |  |  |  |  | Insects | Likely aerial and gleaning |
| **MINIOPTERIDAE** |  |  |  |  |  |  |  |  |  |  |  |
| *Miniopterus tristis / Pipistrellus collinus* | 38 st. cFM |  |  |  |  |  |  | 11 |  | Aerial insects | Hawker |
| *Miniopterus* sp. 1 'medium' | 43 st.cFM | 9 | 11 |  |  | 7 | 2 |  |  | Likely insects | NA |
| *Miniopterus australis* [=*Miniopterus* sp. 2 ‘small’] | 54 st.cFM | 5 | 1 |  |  |  | 39 | 13 | 4 | Flies, ants, moths and wasps | Hawker |
| **MOLOSSIDAE** |  |  |  |  |  |  |  |  |  |  |  |
| *Austronomus kuboriensis* | 13 cFM |  | 5 |  |  |  | 3 |  |  | Beetles | Hawker |
| *Otomops secundus* | 18 cFM |  |  |  |  |  | 1 |  |  | Beetles, aerial insects | Hawker |
| Total species richness | | 7 | 5 | 2 | 1 | 2 | 4 | 2 | 1 |  |  |
| Total relative abudances | | 55 | 37 | 11 | 1 | 29 | 45 | 24 | 4 |  |  |

Zachos, F.E. D. E. Wilson and R. A. Mittermeier (chief editors): Handbook of the Mammals of the World. Vol. 9. Bats.. *Mamm Biol* **100,** 335 (2020). https://doi.org/10.1007/s42991-020-00026-w

**Table S6.** Average increase of abundances of all arthropods in predator exclosures in wet and dry season at the surveyed elevational study sites based on the raw data (a) and based on the estimates predicted by the best model (b). Emmean estimates (below diagonal) and pairwise contrasts (above diagonal) for the treatments irrespective to the season and elevation. Treatments: VER = insectivorous vertebrates excluded, ALL = both insectivorous vertebrates and ants excluded, ANT = ants excluded, CON = control treatment to which the abundances of arthropods in predator exclosures were related.

|  | a) | Abundance increased by %  (Raw data) | | |  | b) Abundance increased by %  (3700 Model) | | |
| --- | --- | --- | --- | --- | --- | --- | --- | --- |
|  |  | VER | ALL | ANT |  | VER | ALL | ANT |
| Wet season | 200 | 39.27 | 36.86 | 3.70 |  | 39.86 | 22.55 | 3.28 |
|  | 700 | 58.72 | 17.76 | 10.29 |  | 32.41 | 22.99 | 6.39 |
|  | 1200 | 58.90 | 38.09 | 20.03 |  | 27.48 | 24.40 | 9.13 |
|  | 1700 | 30.21 | 44.97 | 6.02 |  | 24.60 | 26.89 | 11.77 |
|  | 2200 | 37.88 | 44.69 | 25.68 |  | 23.67 | 30.81 | 14.54 |
|  | 2700 | 92.85 | 74.99 | 16.12 |  | 24.99 | 36.84 | 17.77 |
|  | 3200 | 73.26 |  |  |  | 29.51 |  |  |
|  | 3700 | 77.15 |  |  |  | 39.82 |  |  |
| Dry season | 200 | 54.66 | 28.58 | 7.64 |  | 68.41 | 38.68 | 5.62 |
|  | 700 | 13.96 | 14.02 | -9.24 |  | 48.74 | 34.55 | 9.59 |
|  | 1200 | 33.35 | 34.24 | 14.70 |  | 37.98 | 33.71 | 12.61 |
|  | 1700 | 48.17 | 29.03 | 4.97 |  | 32.21 | 35.20 | 15.40 |
|  | 2200 | -4.67 | 19.14 | -16.95 |  | 30.04 | 39.09 | 18.44 |
|  | 2700 | 25.39 | 38.19 | 14.45 |  | 31.41 | 46.31 | 22.33 |
|  | 3200 | 31.63 |  |  |  | 37.77 |  |  |
|  | 3700 | 10.74 |  |  |  | 54.56 |  |  |
|  | c) |  | CON | ANT | VER | | ALL |  |
|  |  | CON | X | 0.390 | **0.003** | | **0.002** |  |
|  |  | ANT | -3.179 | X | 0.354 | | 0.233 |  |
|  |  | VER | **-6.484** | -3.305 | X | | 0.985 |  |
|  |  | ALL | **-7.204** | -4.024 | -0.719 | | X |  |

**Table S7.** Average increase of herbivory chewing damage in predator exclosures in wet and dry season at the surveyed elevational study sites based on the raw data (a) and based on the estimates predicted by the best model (b). Emmean estimates (below diagonal) and pairwise contrasts (above diagonal) for the treatments irrespective to the season and elevation. Treatments: VER = insectivorous vertebrates excluded, ALL = both insectivorous vertebrates and ants excluded, ANT = ants excluded, CON = control treatment to which the abundances of arthropods in predator exclosures were related.

|  | a) | Increased by % (Raw data) | | |  | Increase by % (3700 Model) | | |
| --- | --- | --- | --- | --- | --- | --- | --- | --- |
|  |  | VER | ALL | ANT |  | VER | ALL | ANT |
| Wet season | 200 | 230.30 | 242.87 | 123.17 |  | 22.13 | 42.38 | 28.15 |
|  | 700 | 70.50 | 66.59 | 5.49 |  | 19.30 | 28.87 | 16.46 |
|  | 1200 | -26.02 | 8.06 | 60.80 |  | 21.00 | 23.79 | 8.19 |
|  | 1700 | -17.75 | -22.66 | -28.54 |  | 27.46 | 26.28 | 2.76 |
|  | 2200 | 85.65 | 59.03 | 140.33 |  | 39.51 | 36.92 | -0.18 |
|  | 2700 | 114.04 | 102.33 | -51.60 |  | 58.71 | 57.87 | -0.81 |
|  | 3200 | 118.11 |  |  |  | 87.74 |  |  |
|  | 3700 | 100.00 |  |  |  | 130.96 |  |  |
| Dry season | 200 | -3.94 | 29.90 | 0.42 |  | 22.35 | 42.88 | 28.45 |
|  | 700 | 7.33 | -14.97 | 40.68 |  | 19.47 | 29.16 | 16.62 |
|  | 1200 | 23.47 | 24.98 | -5.84 |  | 21.16 | 23.98 | 8.27 |
|  | 1700 | 250.05 | 114.17 | 87.01 |  | 27.64 | 26.46 | 2.79 |
|  | 2200 | 35.99 | 72.64 | 36.19 |  | 39.73 | 37.12 | -0.18 |
|  | 2700 | 66.67 | 193.78 | 34.95 |  | 59.01 | 58.15 | -0.81 |
|  | 3200 | 159.82 |  |  |  | 88.13 |  |  |
|  | 3700 | 554.72 |  |  |  | 131.46 |  |  |
|  | c) |  | CON | ANT | VER | | ALL |  |
|  |  | CON | X | 0.743 | **0.001** | | **0.003** |  |
|  |  | ANT | -1.011 | X | **0.062** | | 0.083 |  |
|  |  | VER | **-3.761** | **-2.487** | X | | 1.000 |  |
|  |  | ALL | **-3.447** | -2.373 | 0.001 | | X |  |

**Figure S2.** *Correlations between effect of vertebrate (a, b) and ant (c) exclosure (LRR calculated from raw data, normalized data) on herbivory (a,b,c) caused by chewing arthropods.*

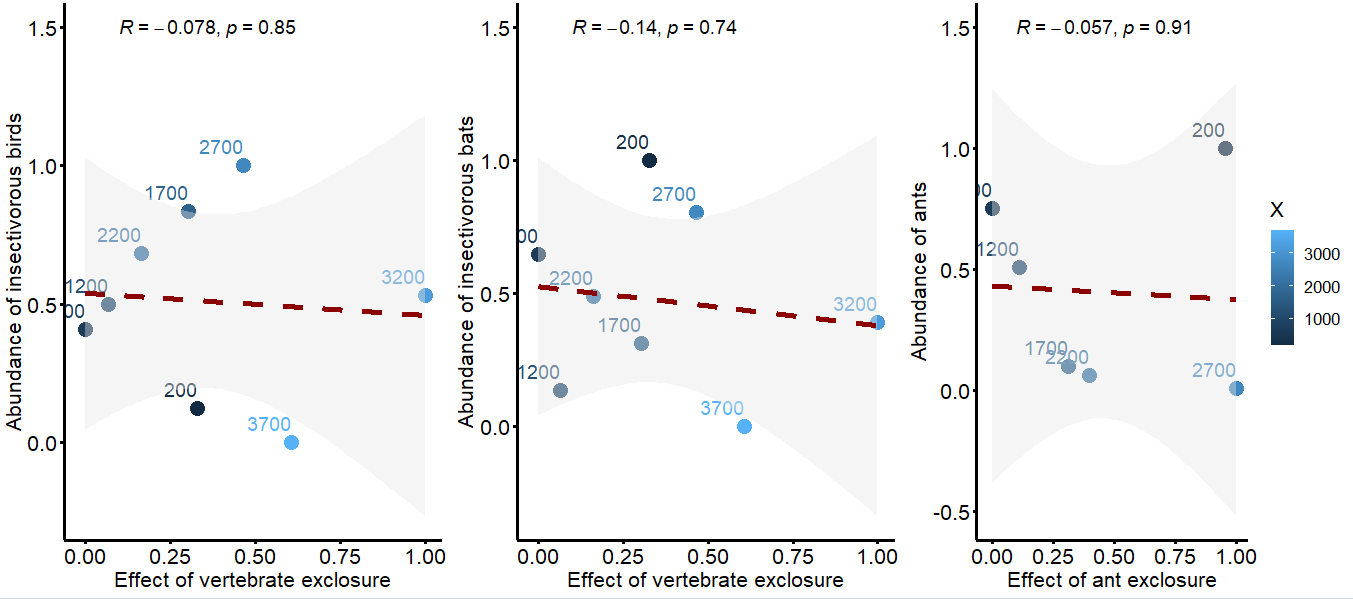
